# Co-allocation Action of Azospirillum brasilense Inoculation and Seed Priming for Improving Germination and Seedling Growth in Aged Grass Seeds

**DOI:** 10.1101/507020

**Authors:** Xv Liu, Zhao Chen, Yani Gao, Qian Liu, Wennan Zhou, Tian Zhao, Wenbo Jiang, Xuewen Cui, Jian Cui, Quanzhen Wang

## Abstract

Germination of seeds during the transportation or after prolonged storage naturally and inevitably decreases because of ageing, but germination potential can be partially restored with seed priming treatments. A novel attempt was made to investigate the effects of combined treatments and to optimize the conditions for naturally aged seeds of tall fescue (*Festuca arundinacea* Schreb.), orchardgrass (*Dactylis glomerata* L.) and Russian wild rye (*Psathyrostachys juncea* (Fisch.) Nevski) using an orthogonal activity level experimental design [factor A: *Azospirillum brasilense* concentration, factor B: three seed priming treatments (H_2_O, MgSO_4_ and H_2_O_2_) and factor C: different priming times]. Multivariate regression model analysis was applied to determine the interactive effects of pairwise factors (A and C) and to optimize experimental conditions. The results showed that the mixed treatments positively affected seed germination and seedling growth. The three seed priming treatments were the dominant factors for germination promotion, whereas the bacterial concentration had the largest effect on seedling growth, especially root elongation. The combined results of all determined attributes showed that *A. brasilense* bio-priming with H_2_O_2_ priming constituted the optimal combination. The optimal bacterial concentration of *A. brasilense* and the time of seed priming were 52.3 × 10^6^ colony forming units (CFU) mL^-1^ and 17.0 h, respectively. We also discussed the physiological mechanisms that could repair aged seeds: the malondialdehyde content and antioxidant enzyme (superoxide dismutase, peroxidase, catalase and ascorbate peroxidase) activities were affected.

## Introduction

The successful completion of seed germination and the establishment of a normal seedling determine the survival and propagation of plant species ^[1]^. However, seeds inevitably and irreversibly deteriorate and age during storage. This deterioration and aging lead to substantial economic losses and decreased genetic diversity ^[1]^. Seed aging can result in reduced germination, seedling abnormalities and even total loss of seed viability, which are correlated with disturbances in various cellular metabolic and biochemical processes, including cell membrane disintegration, mitochondrial dysfunction, enzyme inactivation, Amadori and Maillard reactions, telomere shortening and damage to biological key players such as carbohydrates, DNA and RNA ^[2]^. Currently, the excessive production of reactive oxygen species (ROS; H_2_O_2_, O_2_^-^, etc.) and their attack on lipids and proteins is thought to be the main cause of seed aging ^[2]^. Plants are well equipped to minimize the damaging effects of seed aging via their antioxidant defense system, which consists of superoxide dismutase (SOD), peroxidase (POD) and catalase (CAT), etc. However, it is not always possible for the antioxidant defense system to scavenge excessive ROS by themselves. Seeds always have greater lipid contents than other plant tissues, particularly forage seeds such as those of tall fescue (*Festuca arundinacea* Schreb.) and Russian wild rye (*Psathyrostachys juncea* (Fisch.) Nevski) ^[3, 4]^. Besides, it is difficult to control the ambient temperature, humidity and seed moisture content during storage. These factors can hasten the aging process of seeds ^[5, 6]^. Previous reports have shown that under ambient storage conditions in eastern Serbia in 2009, the reduction in germination of Italian ryegrass (*Lolium multiflorum* L.) and timothy (*Phleum pretense* L.) seeds occurred from 180_th_ and 330_th_ days after harvest, respectively; the germination rates of *Jatropha curcas* L. and lettuce (*Lactuca sativa)* seeds stored for one year decreased by 89.29 and 47.12%, respectively ^[7–9]^. Therefore, exploring effective and pragmatic techniques for seed recovery from aging is urgently warranted for plant germplasm conservation and better agriculture development.

Recent attempts to repair aged seeds through conventional strategies have been met with limited advancement due to the complex nature of seed aging. Behtari and Tilaki reported that the germination percentage of osmoprimed tall fescue seeds stored at 25°C for 1 year decreased significantly, most reducing by 60.1% ^[10]^. In this scenario, the possibility of the application of plant growth-promoting rhizobacteria (PGPR) has emerged as a hot research topic. Various studies have reported that PGPR are capable of alleviating abiotic stress and enhancing growth in plants ^[11, 12]^. *Azospirillum* is versatile and one of the most widely studied genus of PGPR (e.g. *Azotobacter* sp., *Bacillus* sp., *Pseudomonas* sp. and *Herbaspirillum* sp.). It can exert a positive influence on plant growth via complex regulatory networks involved in biological nitrogen fixation, enzyme activation, the production of growth regulatory substances (auxins, gibberellins, polyamines, etc.) and the regulation of the expression of genes in roots ^[13, 14]^.

Seed bio-priming with PGPR can allow enough numbers of bacteria to infect seeds, which is the most promising method for the application of PGPR ^[15]^. Numerous studies have been conducted using *A. brasilense* bio-priming or a consortium of PGPR strains on plants against diverse abiotic stresses, including salinity, osmotic stress and suboptimal temperature ^[16, 17]^. Currently, the fact that the combination of bio-priming with PGPR and another type of seed priming approach (e.g., hydropriming, osmopriming, or redox priming) results in better plant performance is attracting increasing amounts of attention ^[18, 19]^. These combinations are more practical and simpler than co-inoculation of PGPR strains, for which direct antagonism, competition and other factors must be considered. However, the effect of bio-priming via *Azospirillum* on aged seeds has been rarely evaluated, and no reports are available on the repair mechanisms of this genus or optimal combinations with other priming treatments on aged seeds.

Tall fescue, orchardgrass (*Dactylis glomerata* L.) and Russian wildrye are considered to be of great importance among perennial forages due to their high productivity, nutritive value, durability and tolerance to unfavorable conditions. However, poor storability constrains their widespread application ^[20]^. Therefore, the goals of this study were to investigate the effects of *A. brasilense* and various seed priming approaches, including hydropriming, osmopriming and redox priming on the germination and seedling growth of naturally aged forage seeds of tall fescue, orchardgrass and Russian wildrye stored for different years at room temperature and to identify the optimal combination using an orthogonal matrix design and model analysis. Aging-related physiological changes were also examined to better understand the mechanisms of co-treatments on the aged seeds. This study could potentially provide new insights into identifying novel PGPR as elicitors of aging-related repair.

## Materials and Methods

### Bacterial Culture, Seed Materials and Experimental Design

*Azospirillum brasilense* Yu62 was kindly provided by the College of Life Science, Northwest A&F University. The Yu62 culture was prepared on agar-Congo red medium at 37°C for 3 days ^[21]^, a single colony of Yu62 was picked and transferred to nitrogen-free semisolid malate (NFb)liquid medium containing 0.1% NH_4_Cl, and incubated for 48 h at 30°C under vigorous agitation (120 rpm). The cells were harvested by centrifugation (8142 × g, 10 min) and were resuspended in autoclaved 0.12 M phosphate-buffered saline (pH 7.2). The prepared bacterial suspensions were adjusted to 10^8^ colony forming units (CFU) mL^-1^, 10^7^ CFU mL^-1^ and 10^6^ CFU mL^-1^ by adjusting OD values and serially diluting.

Seeds of tall fescue, orchardgrass and Russian wild rye were acquired from the China Agriculture University Grassland Research Station located at the Hexi Corridor in Jiuquan, Gansu Province, China. All seeds were stored for 1 to10 years in paper bags at room temperature in the same laboratory. Six groups of the three grass species, namely, tall fescue 2015 the year harvested, tall fescue 2014, tall fescue 2006, Russian wild rye 2009, orchardgrass 2013 and orchardgrass 2006 that had initial seed moisture contents of 9.3%, 10.1%, 8.2%, 7.9%, 7.7% and 7.9%, respectively, composed the nested experiments. Each group was fixed in an [L9 (3^4^)] orthogonal matrix design for seed priming treatments.

Three factors were selected: the bacterial concentration of *A. brasilense* (factor A); the seed priming approaches viz., hydropriming (autoclaved distilled water), osmopriming (0.37 mol/L MgSO_4_), and redox priming (H_2_O_2_-priming: 0.1% H_2_O_2_) were selected based on numerous previous studies and pre-experiments (factor B) ^[4, 8]^; and priming time (factor C). Each factor was assigned three levels, and the nine treatment combinations of different parameters were laid out in an [L_9_ (3^4^)] orthogonal matrix design replicated thrice (Table 1). Additionally, a control was included without any priming treatment (Table 1).

**Table 1.** The L_9_ (3^4^) orthogonal matrix in the experimental design.

| Treatments | A: the concentration of *A. brasiliense* (CFU mL<sup>-1</sup>) |  | B: seed priming treatment | C : priming time (h) |
| --- | --- | --- | --- | --- |
| | matrix | Code ($X_1$) | matrix | matrix |
| T1 | 1 ($10^6$) | 1 | 1 (hydropriming) | 3 (24) |
| T2 | 2 ($10^7$) | 10 | 1 | 1 (12) |
| T3 | 3 ($10^8$) | 100 | 1 | 2 (18) |
| T4 | 1 | 1 | 2 (MgSO<sub>4</sub>-priming) | 2 |
| T5 | 2 | 10 | 2 | 3 |
| T6 | 3 | 100 | 2 | 1 |
| T7 | 1 | 1 | 3 (H<sub>2</sub>O<sub>2</sub>-priming) | 1 |
| T8 | 2 | 10 | 3 | 2 |
| T9 | 3 | 100 | 3 | 3 |

### Seed Treatments and Germination Tests

Seeds were surface-sterilized with 2% NaOCl solution for 5 min followed by 70% ethanol for 10 min, after which the seeds were rinsed five times with sterile distilled water. According to the orthogonal design [L_9_ (3^4^)], each group of seeds that were coated with gauze were primed at ambient temperature. Inoculation was performed by soaking the sterilized seeds in prepared bacterial suspensions or only sterile phosphate-buffered saline for control seeds, for 2 h ^[15]^. After hydropriming, osmopriming, or redox priming followed by *A. brasilense* inoculation, the primed seeds were air-dried to reduce their moisture content. Three replicates were used for each treatment and 50 healthy seeds from each replication were evenly spaced on two layers of moistened Whatman No. 1 filter paper in each 150-mm Petri dish. Equal volumes of distilled water were added daily. Petri dishes were placed in a climate incubator (RQH-250, Jing Hong Laboratory Instrument Co., Ltd, Shanghai, China) that maintained an alternating diurnal regimen of 16 h of light at 25 ± 0.5°C and 8 h of dark at 20 ± 0.5°C at 85% relative humidity for 28 days. Seed germination was recorded daily, and the seed was considered germinated when its radicle extended at least 0.5 mm. The shoot and root lengths of 10 randomly selected seedlings from each replication were measured in accordance with the rules recommended by the International Seed Testing Association (ISTA) at 28_th_ day after sowing (DAS). When the total number of seedlings was fewer than 10, all seedlings were measured. To more precisely reflect the vigor of seeds, the germination index (Gi) was estimated as follows:

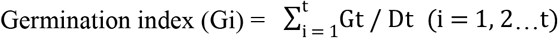

where Gt is the is the number of germinated seeds within a day, Dt is the corresponding number of germination days and t is the number of total germination period (28 days).

The seed vigour index (SVi) was determined using the following formula:

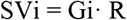

where R is length of root.

### Determination of Physiological Parameters in Aged Grass Seeds

Lipid peroxidation was assessed in terms of the malondialdehyde (MDA) content in fresh seedlings according to the thiobarbituric acid protocol ^[22]^. Each group of sample containing 0.5 g crumbled seedlings was mixed with 5 mL trichloroacetic acid (TCA; 0.5%) and was centrifuged at 10,000 × g for 25 min. For every 1 mL of the aliquot of the supernatant, 4 mL of 20% TCA containing 0.5% thiobarbituric acid was added. The mixture was heated at 95 °C for 30 min and then the tubes were cooled immediately in an ice-bath. After the tube was centrifuged at 10 000 × g for 10 min, the absorbance of supernatant was measured at 450, 532, and 600 nm. The MDA content was expressed as μmol·g^-1^ FW.

The activities of the antioxidant enzymes SOD, POD, CAT and ascorbate peroxidase (APX) in fresh seedlings were determined. The extraction of antioxidant enzymes was carried out on seedling samples in accordance with the procedures described by Zhang and Nan, with minor modifications ^[23]^. Each group of seedlings (0.5 g) was homogenized in 5 mL of 50 mM phosphate buffer (pH 7.8 for SOD and POD containing 1% (w/v) polyvinylpyrrolidone, pH 7 for CAT and APX containing 0.1 mM Na_2_EDTA). The homogenate was centrifuged at 10000 × g for 15 min at 4°C, after which the supernatant was collected for the determination of enzyme activities. The activities of SOD, POD, CAT and APX were analyzed as reported by Zhang and Nan ^[23]^.

### Statistical Analysis

The data were statistically analyzed by means of variance analysis (ANOVA) using the software SPSS version 20.0 and were compared using the Duncan test (*P* < 0.05).

The below-mentioned analyses and graphical procedures were performed using SAS version 8.2. The germination percentage (GP) was simulated using the following logistic model ^[24, 25]^: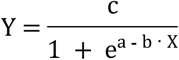 where a, b, and c are constants and X represents the germination days. The models were significant at *P* < 0.05.

For the generic results, the bacterial concentration of *A. brasilense* and seed priming time (factors A and C) were denoted by X_1_ and X_3_. The dependent variables were denoted by Y_i_, which included GP, Gi, SVi, root length, shoot length, seedling length, the shoot/root ratio (the index of seed vigor during aging), MDA content, and the activity of SOD, POD, CAT and APX. These dependent variables were approached and analyzed via two-variable (X_1_ and X_3_) quadratic regression models described as follows:

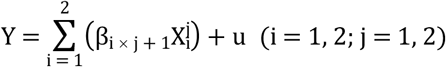

where β is a constant. The critical values of X_1_ and X_3_ were obtained from the quadratic models that were significant at *P* < 0.05.

## Results

### The effect of co-treatment on the vigor of aged seeds

None of the seeds of tall fescue 2006 germinated in the whole nested experiment, only 2-4% of seeds germinated in treatments 1-4 (T1-T4) and T7 for orchardgrass 2006 (data not shown). The models regarding germination dynamics (Fig 1) showed that the germination of T1-T5, and T8 in tall fescue 2015 occurred earlier than it did for the control (CK), and the greatest germination percentage (28 DAS) occurred in T1 (98.7%). The germination from all treatments occurred earlier than it did from the controls for both orchardgrass 2013 and Russian wildrye 2009. The other 8 treatments increased germination of orchardgrass 2013 except for T3, the germination percentages from T2 and T6 (92.0%) were the highest. The germination percentages of all treatments were higher than that of the CK, the highest germination percentage was in T1 (62.7%) among them in Russian wildrye 2009. Whereas, the co-treatments had a negative effect on germination of tall fescue 2014 except for T7 (72.0%) compared with the CK. But the germination started earlier in T2 and T7. The lowest germination percentages of tall fescue 2015, tall fescue 2014 and orchardgrass 2013 occurred in T9, T3 and T3, respectively. While the lowest germination percentage of Russian wildrye 2009 occurred in T9, which increased by 2.7 times compared with that of the CK.

**Fig 1.**
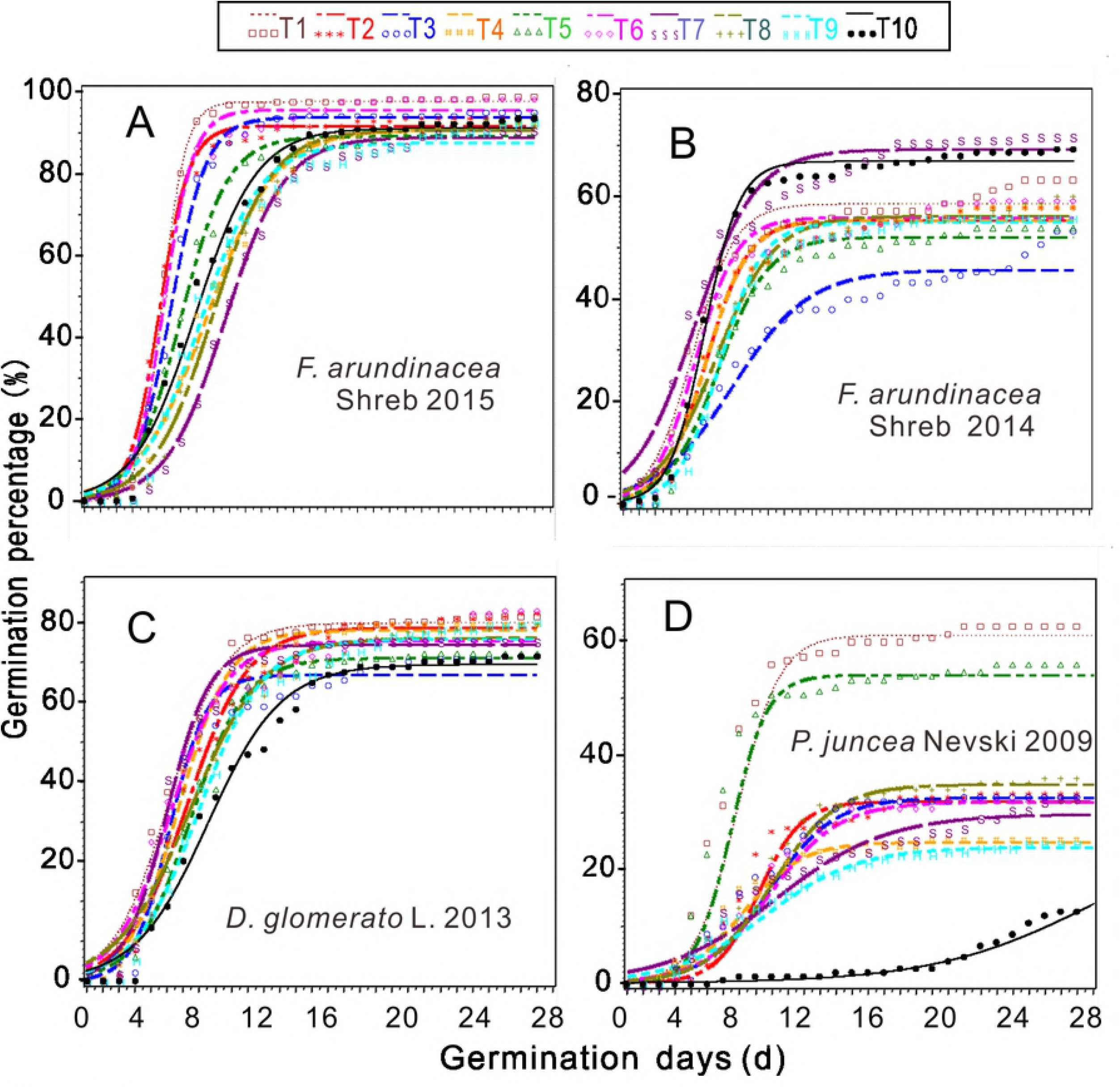
The curves of logistic models for germination percentage under the combined treatments for tall fescue 2015 (A), 2014 (B), orchardgrass 2013 (C) and Russian wild rye 2009 (D). From T1 to T9 are the treatments, and T10 is the control.

The range analyses revealed that factor C (seed priming time) exhibited the largest effect on germination percentage among three factors, with the exception that factor B (seed priming approaches) had the highest range value (R value; 4.0) for tall fescue 2015 (Table 2). And the most effective factors on the Gi and SVi of tall fescue 2015, orchardgrass 2013 and Russian wildrye 2009 were factors B, A (the bacterial concentration) and C, individually, which were the same for every species. The most important factors on the Gi and SVi of tall fescue 2014 were factors C and B, which were different. The analysis of variance indicated that the germination percentage was significantly (*P* < 0.05) affected by both grass species and factor C, and their interactive effects were also significant (*P* < 0.05) both in pairwise factors and among three factors, with the exceptions of the grass species × factor A and grass species × factor B interaction (Table 3). Both factors B and C significantly (*P* < 0.01) affected the Gi and SVi in different-aged seeds, also their interactive effect was significant (*P*<0.05).

**Table 2.**
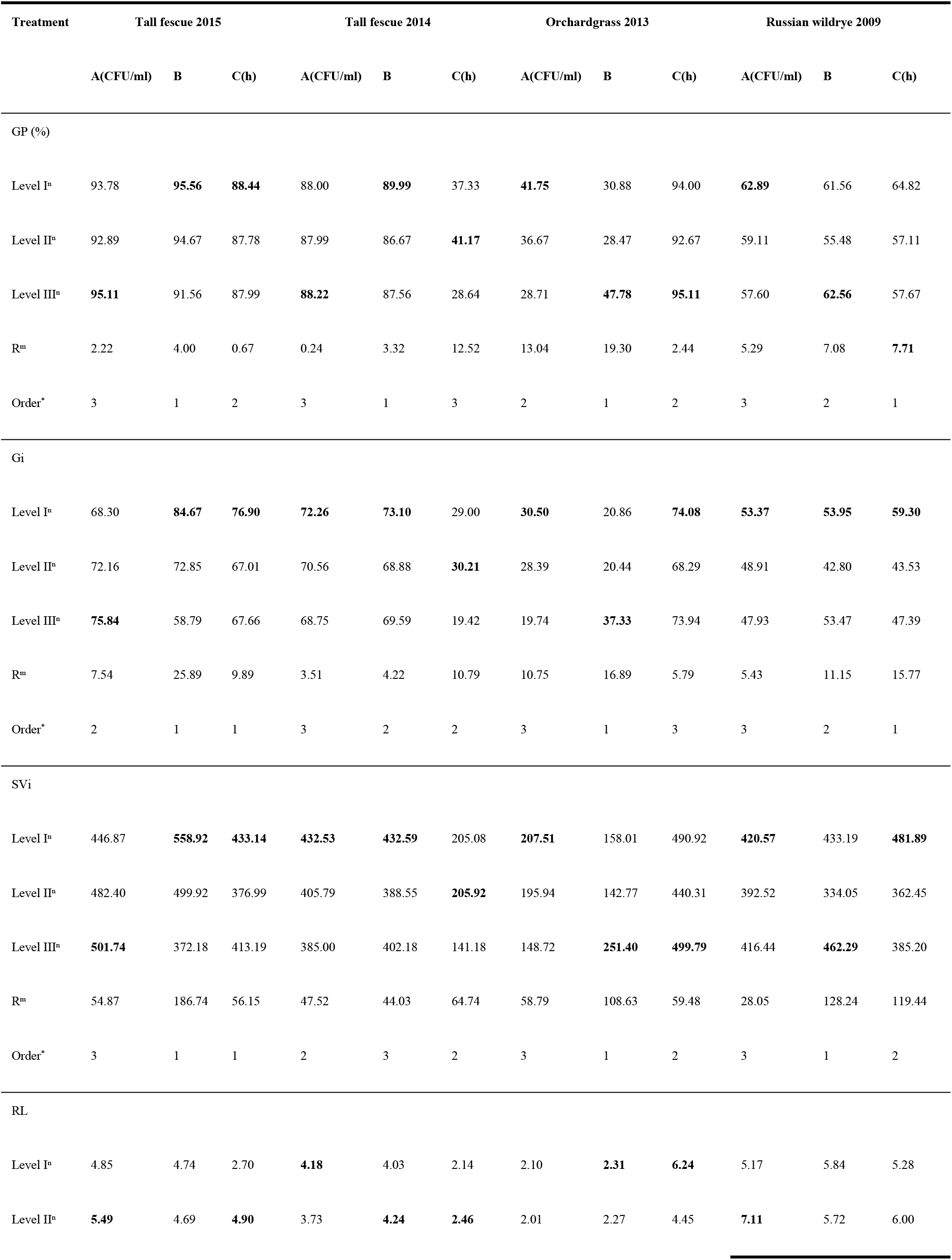
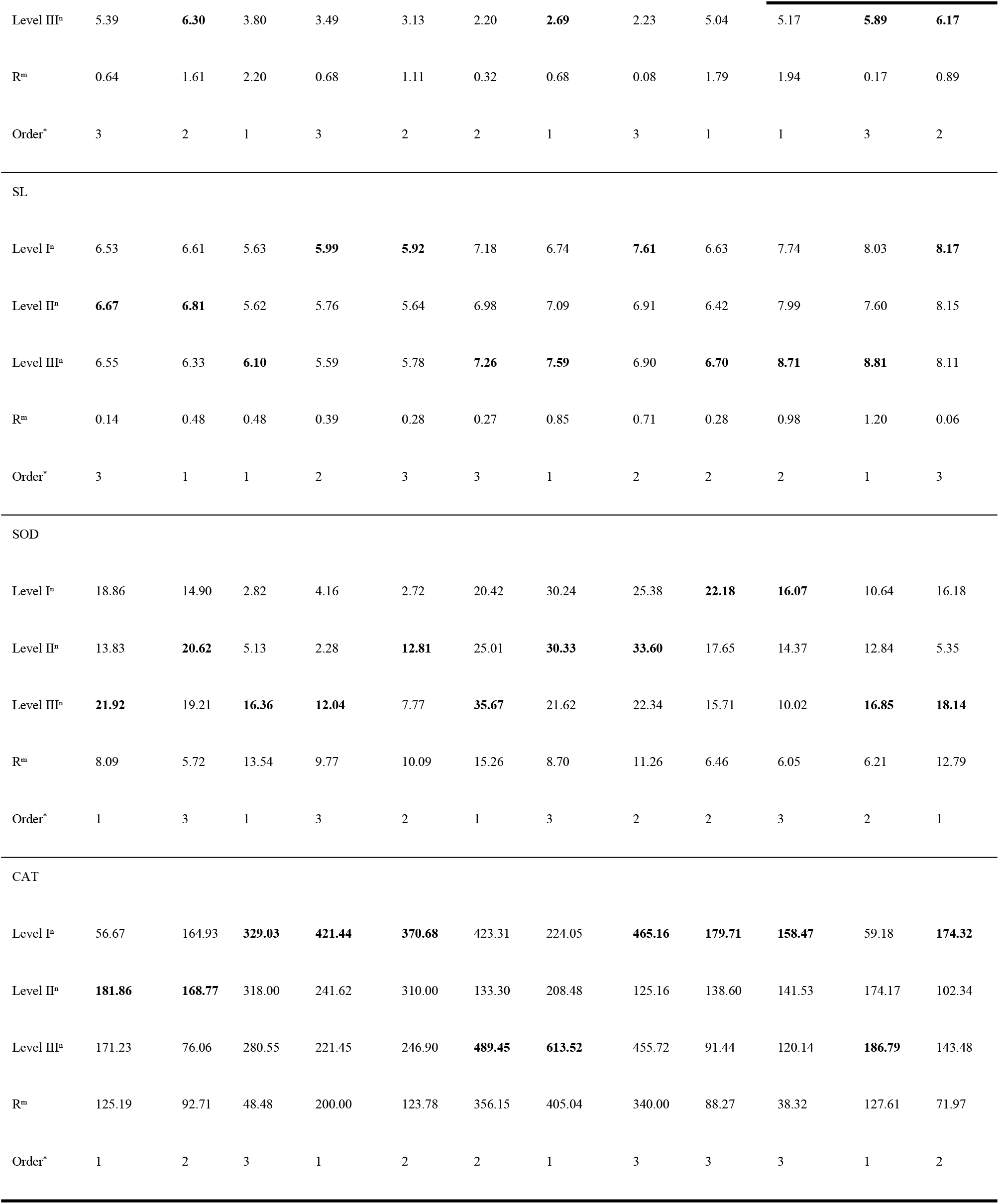
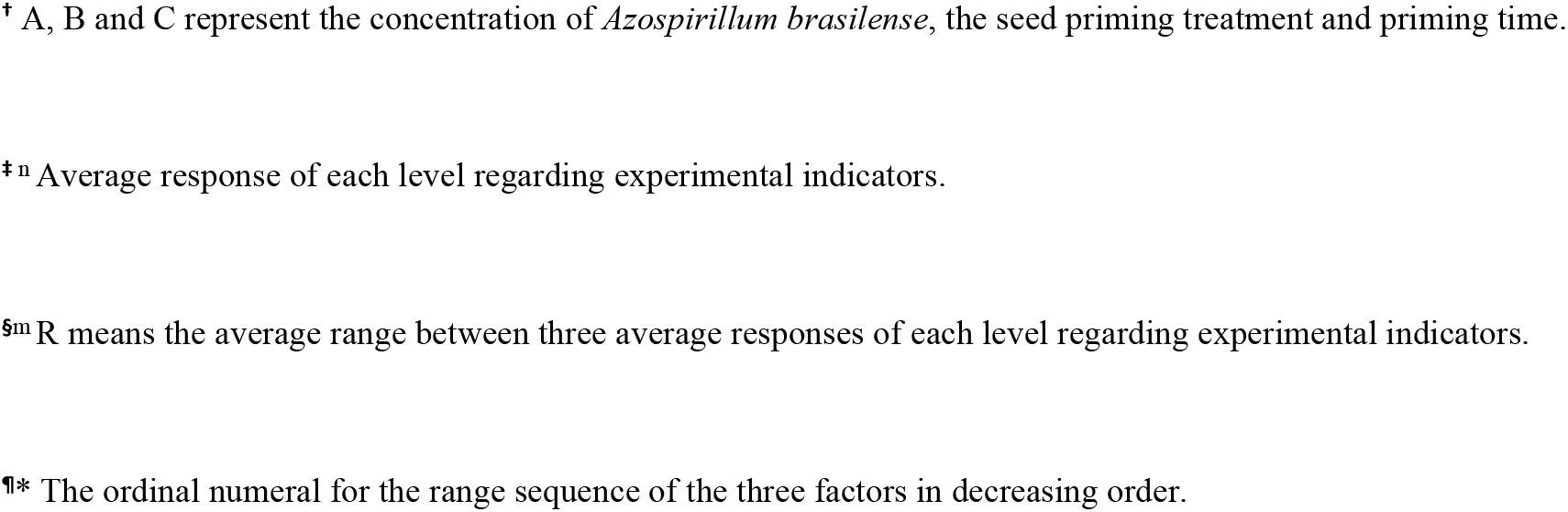
Average responses of each level and the range analyses of germination percentage (GP), germination index (Gi), seed vigor index (SVi), root (RL) and shoot (RL) length in tall fescue 2015, tall fescue 2014, orchardgrass 2013 and Russian wild rye 2009.

| Treatment | Tall fescue 2015 |  |  | Tall fescue 2014 |  |  | Orchardgrass 2013 |  |  | Russian wildrye 2009 |  |  |
| --- | --- | --- | --- | --- | --- | --- | --- | --- | --- | --- | --- | --- |
|  | A(CFU/ml) | B | C(h) | A(CFU/ml) | B | C(h) | A(CFU/ml) | B | C(h) | A(CFU/ml) | B | C(h) |
| GP (%) |  |  |  |  |  |  |  |  |  |  |  |  |
| Level I<sup>a</sup> | 93.78 | **95.56** | **88.44** | 88.00 | **89.99** | 37.33 | **41.75** | 30.88 | 94.00 | **62.89** | 61.56 | 64.82 |
| Level II<sup>a</sup> | 92.89 | 94.67 | 87.78 | 87.99 | 86.67 | **41.17** | 36.67 | 28.47 | 92.67 | 59.11 | 55.48 | 57.11 |
| Level III<sup>a</sup> | **95.11** | 91.56 | 87.99 | **88.22** | 87.56 | 28.64 | 28.71 | **47.78** | **95.11** | 57.60 | **62.56** | 57.67 |
| R<sup>m</sup> | 2.22 | 4.00 | 0.67 | 0.24 | 3.32 | 12.52 | 13.04 | 19.30 | 2.44 | 5.29 | 7.08 | **7.71** |
| Order<sup>a</sup> | 3 | 1 | 2 | 3 | 1 | 3 | 2 | 1 | 2 | 3 | 2 | 1 |
| Gi |  |  |  |  |  |  |  |  |  |  |  |  |
| Level I<sup>a</sup> | 68.30 | **84.67** | **76.90** | **72.26** | **73.10** | 29.00 | **30.50** | 20.86 | **74.08** | **53.37** | **53.95** | **59.30** |
| Level II<sup>a</sup> | 72.16 | 72.85 | 67.01 | 70.56 | 68.88 | **30.21** | 28.39 | 20.44 | 68.29 | 48.91 | 42.80 | 43.53 |
| Level III<sup>a</sup> | **75.84** | 58.79 | 67.66 | 68.75 | 69.59 | 19.42 | 19.74 | **37.33** | 73.94 | 47.93 | 53.47 | 47.39 |
| R<sup>m</sup> | 7.54 | 25.89 | 9.89 | 3.51 | 4.22 | 10.79 | 10.75 | 16.89 | 5.79 | 5.43 | 11.15 | 15.77 |
| Order<sup>a</sup> | 2 | 1 | 1 | 3 | 2 | 2 | 3 | 1 | 3 | 3 | 2 | 1 |
| SVi |  |  |  |  |  |  |  |  |  |  |  |  |
| Level I<sup>a</sup> | 446.87 | **558.92** | **433.14** | **432.53** | **432.59** | 205.08 | **207.51** | 158.01 | 490.92 | **420.57** | 433.19 | **481.89** |
| Level II<sup>a</sup> | 482.40 | 499.92 | 376.99 | 405.79 | 388.55 | **205.92** | 195.94 | 142.77 | 440.31 | 392.52 | 334.05 | 362.45 |
| Level III<sup>a</sup> | **501.74** | 372.18 | 413.19 | 385.00 | 402.18 | 141.18 | 148.72 | **251.40** | **499.79** | 416.44 | **462.29** | 385.20 |
| R<sup>m</sup> | 54.87 | 186.74 | 56.15 | 47.52 | 44.03 | 64.74 | 58.79 | 108.63 | 59.48 | 28.05 | 128.24 | 119.44 |
| Order<sup>a</sup> | 3 | 1 | 1 | 2 | 3 | 2 | 3 | 1 | 2 | 3 | 1 | 2 |
| RL |  |  |  |  |  |  |  |  |  |  |  |  |
| Level I<sup>a</sup> | 4.85 | 4.74 | 2.70 | **4.18** | 4.03 | 2.14 | 2.10 | **2.31** | **6.24** | 5.17 | 5.84 | 5.28 |
| Level II<sup>a</sup> | **5.49** | 4.69 | **4.90** | 3.73 | **4.24** | **2.46** | 2.01 | 2.27 | 4.45 | **7.11** | 5.72 | 6.00 |
| Level III<sup>a</sup> | 5.39 | **6.30** | 3.80 | 3.49 | 3.13 | 2.20 | **2.69** | 2.23 | 5.04 | 5.17 | **5.89** | **6.17** |
| R<sup>m</sup> | 0.64 | 1.61 | 2.20 | 0.68 | 1.11 | 0.32 | 0.68 | 0.08 | 1.79 | 1.94 | 0.17 | 0.89 |
| Order<sup>a</sup> | 3 | 2 | 1 | 3 | 2 | 2 | 1 | 3 | 1 | 1 | 3 | 2 |
| SL |  |  |  |  |  |  |  |  |  |  |  |  |
| Level I<sup>a</sup> | 6.53 | 6.61 | 5.63 | **5.99** | **5.92** | 7.18 | 6.74 | **7.61** | 6.63 | 7.74 | 8.03 | **8.17** |
| Level II<sup>a</sup> | **6.67** | **6.81** | 5.62 | 5.76 | 5.64 | 6.98 | 7.09 | 6.91 | 6.42 | 7.99 | 7.60 | 8.15 |
| Level III<sup>a</sup> | 6.55 | 6.33 | **6.10** | 5.59 | 5.78 | **7.26** | **7.59** | 6.90 | **6.70** | **8.71** | **8.81** | 8.11 |
| R<sup>m</sup> | 0.14 | 0.48 | 0.48 | 0.39 | 0.28 | 0.27 | 0.85 | 0.71 | 0.28 | 0.98 | 1.20 | 0.06 |
| Order<sup>a</sup> | 3 | 1 | 1 | 2 | 3 | 3 | 1 | 2 | 2 | 2 | 1 | 3 |
| SOD |  |  |  |  |  |  |  |  |  |  |  |  |
| Level I<sup>a</sup> | 18.86 | 14.90 | 2.82 | 4.16 | 2.72 | 20.42 | 30.24 | 25.38 | **22.18** | **16.07** | 10.64 | 16.18 |
| Level II<sup>a</sup> | 13.83 | **20.62** | 5.13 | 2.28 | **12.81** | 25.01 | **30.33** | **33.60** | 17.65 | 14.37 | 12.84 | 5.35 |
| Level III<sup>a</sup> | **21.92** | 19.21 | **16.36** | **12.04** | 7.77 | **35.67** | 21.62 | 22.34 | 15.71 | 10.02 | **16.85** | **18.14** |
| R<sup>m</sup> | 8.09 | 5.72 | 13.54 | 9.77 | 10.09 | 15.26 | 8.70 | 11.26 | 6.46 | 6.05 | 6.21 | 12.79 |
| Order<sup>a</sup> | 1 | 3 | 1 | 3 | 2 | 1 | 3 | 2 | 2 | 3 | 2 | 1 |
| CAT |  |  |  |  |  |  |  |  |  |  |  |  |
| Level I<sup>a</sup> | 56.67 | 164.93 | **329.03** | **421.44** | **370.68** | 423.31 | 224.05 | **465.16** | **179.71** | **158.47** | 59.18 | **174.32** |
| Level II<sup>a</sup> | **181.86** | **168.77** | 318.00 | 241.62 | 310.00 | 133.30 | 208.48 | 125.16 | 138.60 | 141.53 | 174.17 | 102.34 |
| Level III<sup>a</sup> | 171.23 | 76.06 | 280.55 | 221.45 | 246.90 | **489.45** | **613.52** | 455.72 | 91.44 | 120.14 | **186.79** | 143.48 |
| R<sup>m</sup> | 125.19 | 92.71 | 48.48 | 200.00 | 123.78 | 356.15 | 405.04 | 340.00 | 88.27 | 38.32 | 127.61 | 71.97 |
| Order<sup>a</sup> | 1 | 2 | 3 | 1 | 2 | 2 | 1 | 3 | 3 | 3 | 1 | 2 |

<sup>†</sup> A, B and C represent the concentration of *Azospirillum brasilense*, the seed priming treatment and priming time.
<sup>‡n</sup> Average response of each level regarding experimental indicators.
<sup>§m</sup> R means the average range between three average responses of each level regarding experimental indicators.
<sup>¶\*</sup> The ordinal numeral for the range sequence of the three factors in decreasing order.

**Table 3.**
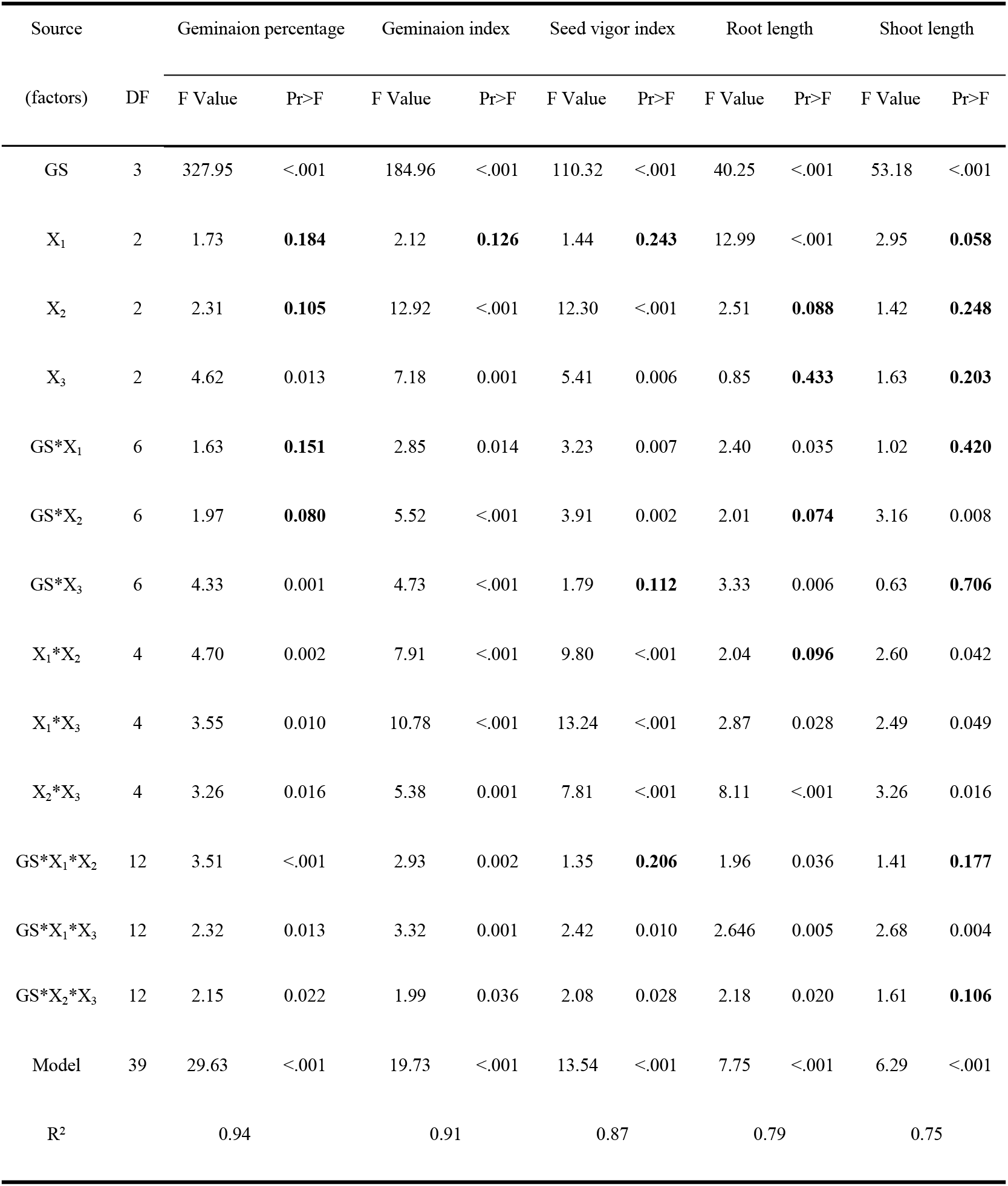
Variance analyses for the model, the effects of each experimental factors and their interaction effects on germination percentage, gemination index, seed vigor index, root, and shoot length. Columns A, X_1_ X_2_ and X_3_ represent grass species, bacteria concentration, seed priming and priming duration, respectively.

| Source<br>(factors) | DF | Geminaion percentage |  | Geminaion index |  | Seed vigor index |  | Root length |  | Shoot length |  |
| --- | --- | --- | --- | --- | --- | --- | --- | --- | --- | --- | --- |
|  |  | F Value | Pr>F | F Value | Pr>F | F Value | Pr>F | F Value | Pr>F | F Value | Pr>F |
| GS | 3 | 327.95 | <.001 | 184.96 | <.001 | 110.32 | <.001 | 40.25 | <.001 | 53.18 | <.001 |
| $X_1$ | 2 | 1.73 | **0.184** | 2.12 | **0.126** | 1.44 | **0.243** | 12.99 | <.001 | 2.95 | **0.058** |
| $X_2$ | 2 | 2.31 | **0.105** | 12.92 | <.001 | 12.30 | <.001 | 2.51 | **0.088** | 1.42 | **0.248** |
| $X_3$ | 2 | 4.62 | 0.013 | 7.18 | 0.001 | 5.41 | 0.006 | 0.85 | **0.433** | 1.63 | **0.203** |
| GS*$X_1$ | 6 | 1.63 | **0.151** | 2.85 | 0.014 | 3.23 | 0.007 | 2.40 | 0.035 | 1.02 | **0.420** |
| GS*$X_2$ | 6 | 1.97 | **0.080** | 5.52 | <.001 | 3.91 | 0.002 | 2.01 | **0.074** | 3.16 | 0.008 |
| GS*$X_3$ | 6 | 4.33 | 0.001 | 4.73 | <.001 | 1.79 | **0.112** | 3.33 | 0.006 | 0.63 | **0.706** |
| $X_1$*$X_2$ | 4 | 4.70 | 0.002 | 7.91 | <.001 | 9.80 | <.001 | 2.04 | **0.096** | 2.60 | 0.042 |
| $X_1$*$X_3$ | 4 | 3.55 | 0.010 | 10.78 | <.001 | 13.24 | <.001 | 2.87 | 0.028 | 2.49 | 0.049 |
| $X_2$*$X_3$ | 4 | 3.26 | 0.016 | 5.38 | 0.001 | 7.81 | <.001 | 8.11 | <.001 | 3.26 | 0.016 |
| GS*$X_1$*$X_2$ | 12 | 3.51 | <.001 | 2.93 | 0.002 | 1.35 | **0.206** | 1.96 | 0.036 | 1.41 | **0.177** |
| GS*$X_1$*$X_3$ | 12 | 2.32 | 0.013 | 3.32 | 0.001 | 2.42 | 0.010 | 2.646 | 0.005 | 2.68 | 0.004 |
| GS*X<sub>2</sub>*X<sub>3</sub> | 12 | 2.15 | 0.022 | 1.99 | 0.036 | 2.08 | 0.028 | 2.18 | 0.020 | 1.61 | **0.106** |
| Model | 39 | 29.63 | <.001 | 19.73 | <.001 | 13.54 | <.001 | 7.75 | <.001 | 6.29 | <.001 |
| R<sup>2</sup> |  | 0.94 |  | 0.91 |  | 0.87 |  | 0.79 |  | 0.75 |  |

### Seedling growth

Compared with the CK, T7 resulted in the biggest increase in root length, and the shoot length increased in T5 and T6 in tall fescue 2015 group (Fig 2). The root length in T8 was the longest significantly, and the shoot lengths in both T8 and T9 were markedly improved in tall fescue 2014. In group of orchardgrass 2013, the root and shoot lengths were increased by T3 and T6, respectively. While in Russian wildrye 2009, the changes of root elongation was not significant. The root length was the largest in T8, the shoot length in T6 was significantly higher than that of the CK.

**Figure 2.**
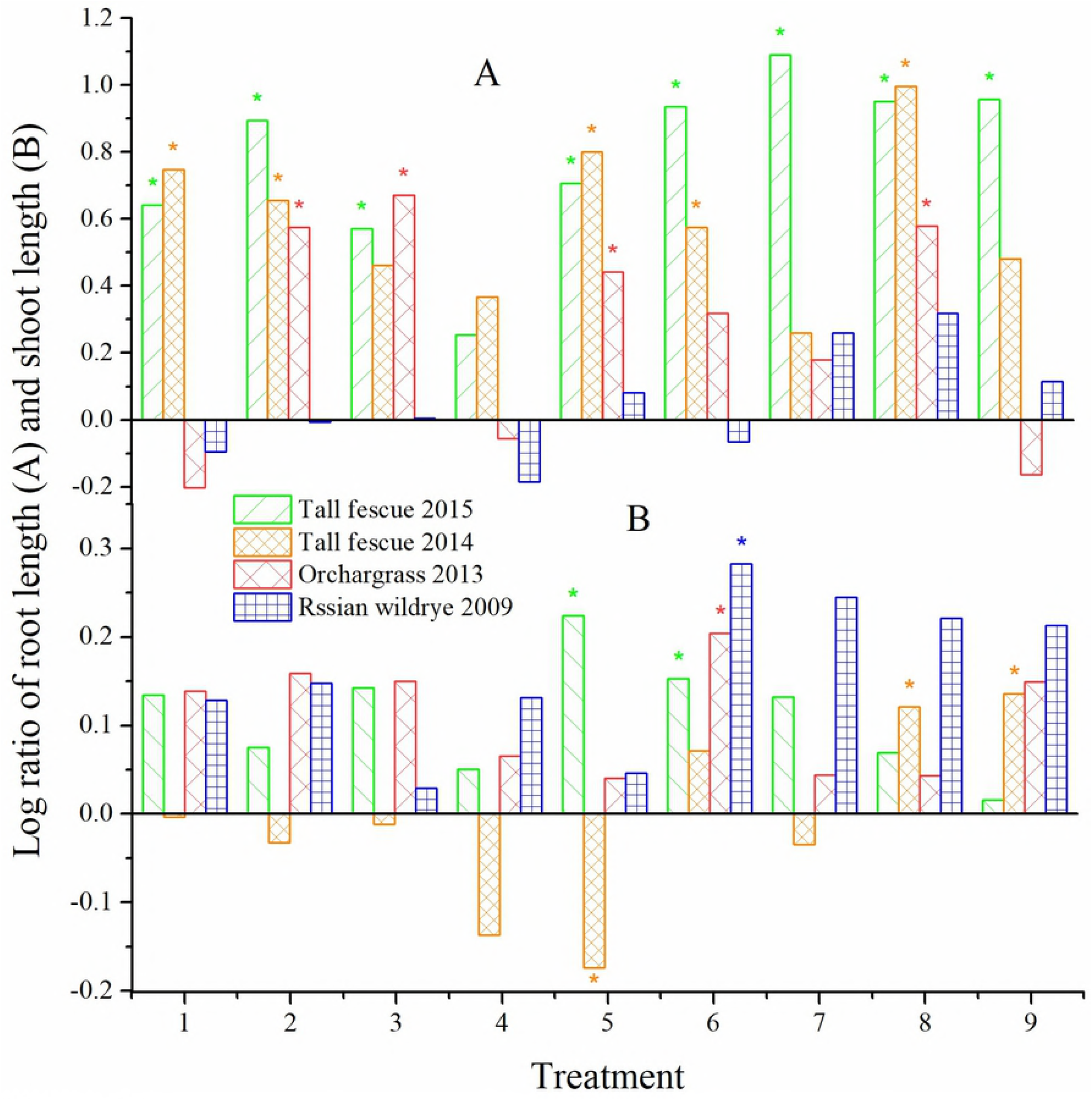
Log ratio of root length (A) and shoot length (B). * indicates the difference was significant between the treatment and the control at *P* < 0.05.

Factor A markedly affected the root length, and the effect of factor A was the largest for tall fescue 2014 and orchardgrass 2013 (Tables 2 and 3). The interactive effects were also significant (*P* < 0.01) on root length, except for the factors A × B. Although factors A, B and C did not significantly influence the shoot length, their pairwise interactive effects were significant (*P* < 0.05; Table 3).

### Characterization of physiological responses

The MDA content of CK in Russian wildrye 2009 was significantly (*P* < 0.05) higher than those of CKs in tall fescue 2015, tall fescue 2014 and orchardgrass 2013. And the SOD activity of CK in Russian wildrye 2009 was noticeably lower than those in the three other controls. These results indicated that the seed of Russian wildrye 2009 was the most severely damaged by aging (Fig 3A; S1 Table). In all treatments, the MDA content sharply decreased relative to that of the CKs in Russian wildrye 2009 and tall fescue 2015. The lowest MDA content in tall fescue 2014 and orchardgrass 2013 occurred in T6 and T7 (Fig 3A). The changing trends regarding the MDA content and SOD activity were opposite in Russian wildrye 2009 and tall fescue 2014 (T1, T4, T5 and CK). The highest activities of SOD and CAT were observed in T6 for tall fescue 2015 and in T7 for tall fescue 2014. While the highest SOD and CAT activities were in T3 and T2 for orchardgrass 2013, and were in T8 and T9 for Russian wildrye 2009, respectively (Fig 3A, B and D). The highest POD values for tall fescue 2015 and Russian wildrye 2009 were found in T3 and T2, separately (Fig 3C).

**Fig 3.**
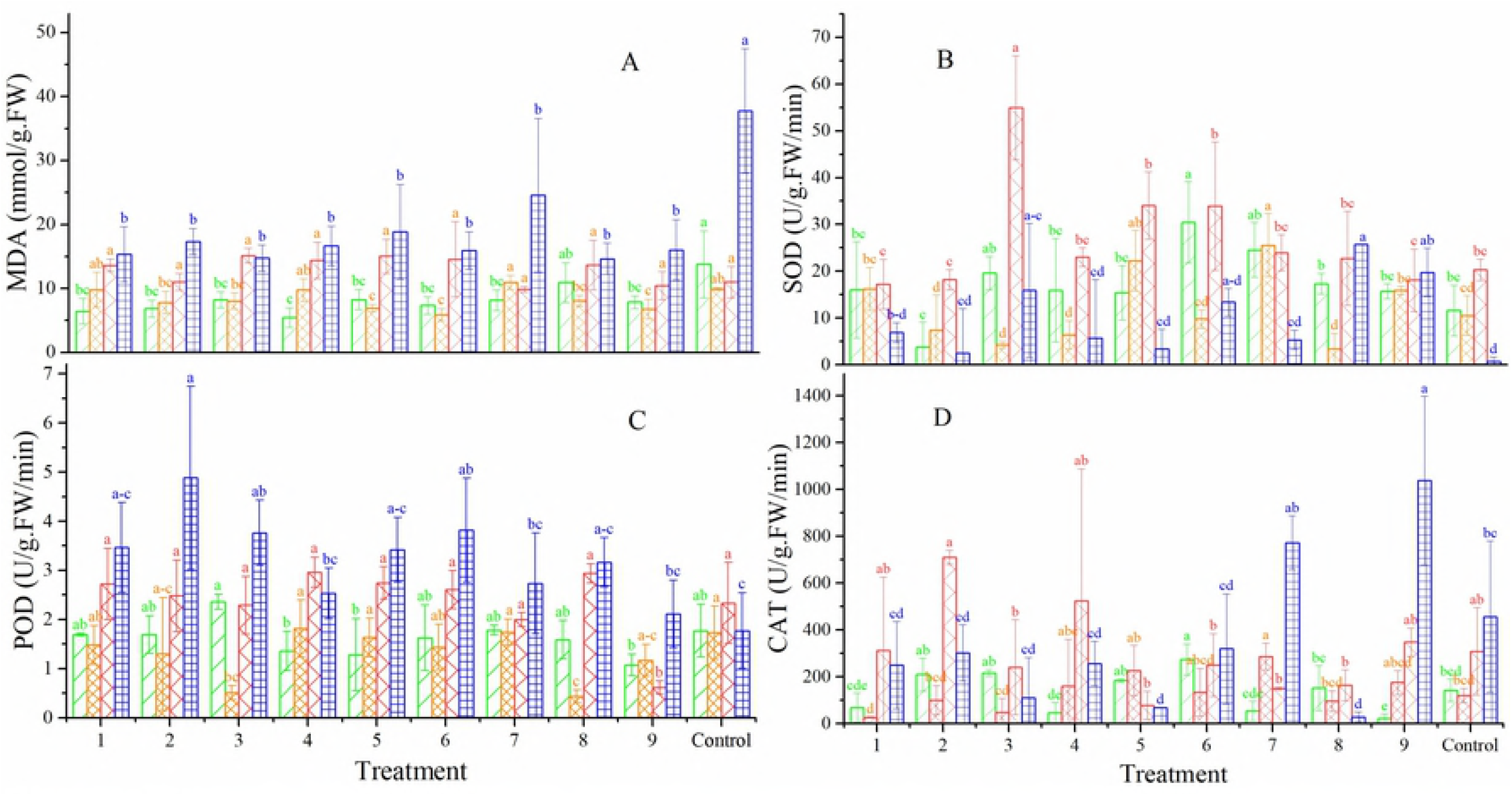
MDA content (A), activity of SOD (B), POD (C) and CAT (D) of tall fescue (2015 and 2014), orchardgrass (2013) and Russian wildrye (2009). Data are presented as mean ± SD. The different lowercase letters indicate significant differences at *P* < 0.05 within the years among the treatments.

Factor A and the interactive effects significantly (*P* < 0.05) influenced SOD activity, and factor A was the most important factor for those of tall fescue 2015, orchardgrass 2013 and Russian wildrye 2009 (Tables 2 and 4). The experimental factors did not significantly affect the MDA content, except seed species (Table 4). The experimental factors markedly affected the POD and CAT activities with the exceptions of factor A, A × B, B × C and grass species ×A × B on the POD activity as well as factors A and B on the CAT activity.

**Table 4.** Variance analyses for the model, the effects of each experimental factors and their interaction effects on MDA, POD, SOD and CAT.

| Source |  | MDA |  | POD |  | SOD |  | CAT |  |
| --- | --- | --- | --- | --- | --- | --- | --- | --- | --- |
| (factors) | DF | F Value | Pr>F | F Value | Pr>F | F Value | Pr>F | F Value | Pr>F |
| GS | 3 | 72.064 | <.001 | 36.214 | <.001 | 36.349 | <.001 | 16.816 | <.001 |
| X<sub>1</sub> | 2 | 0.947 | **0.392** | 2.871 | **0.063** | 14.803 | <.001 | 2.410 | **0.096** |
| X<sub>2</sub> | 2 | 0.277 | **0.759** | 9.357 | <.001 | 2.649 | **0.077** | 1.588 | **0.211** |
| X<sub>3</sub> | 2 | 0.144 | **0.866** | 3.445 | 0.037 | 1.474 | **0.235** | 4.796 | 0.011 |
| GS*X<sub>1</sub> | 6 | 1.259 | **0.286** | 3.294 | 0.006 | 6.447 | <.001 | 4.941 | <.001 |
| GS*X<sub>2</sub> | 6 | 1.302 | **0.266** | 3.333 | 0.006 | 3.063 | 0.009 | 7.143 | <.001 |
| GS*X<sub>3</sub> | 6 | 1.330 | **0.254** | 2.299 | 0.042 | 8.441 | <.001 | 3.490 | 0.004 |
| X<sub>1</sub>*X<sub>2</sub> | 4 | 0.926 | **0.453** | 2.482 | **0.0502** | 6.727 | <.001 | 7.216 | <.001 |
| X<sub>1</sub>*X<sub>3</sub> | 4 | 0.993 | **0.416** | 5.438 | 0.001 | 7.314 | <.001 | 5.612 | <.001 |
| X<sub>2</sub>*X<sub>3</sub> | 4 | 1.328 | **0.267** | 2.195 | **0.077** | 13.392 | <.001 | 6.023 | <.001 |
| GS*X<sub>1</sub>*X<sub>2</sub> | 12 | 1.103 | **0.369** | 1.771 | **0.067** | 8.083 | <.001 | 4.525 | <.001 |
| GS*X<sub>1</sub>*X<sub>3</sub> | 12 | 1.089 | **0.381** | 2.288 | 0.015 | 5.394 | <.001 | 6.351 | <.001 |
| GS*X<sub>2</sub>*X<sub>3</sub> | 12 | 1.067 | **0.398** | 2.268 | 0.016 | 7.086 | <.001 | 5.250 | <.001 |
| Model | 39 | 8.009 | <.0001 | 6.827 | <.001 | 8.838 | <.001 | 5.591 | <.001 |
| R<sup>2</sup> |  | 0.796 |  | 0.769 |  | 0.812 |  | 0.732 |  |

### Model analysis for interactive effects and optimized conditions

Given the interactive effects of factors A and C and exploring optimized conditions, regression models were constructed. As shown in Figure 4, the root and seedling length in the four groups of aged seeds showed via three-dimensional parabolic trends that there were maximum values with increase in factor A. The root and seedling length were sharply increased when the bacterial concentration was in the range of 30.0-75.0 × 10^6^ CFU mL^-1^. The SVi also had maximum values with a bacterial concentration of 30.0-75.0 × 10^6^ CFU mL^-1^ when priming duration was less than 24.0 h, although the SVi of orchardgrass 2013 exhibited a completely opposite trend (Fig 4E-H). The Gi of Tf 2015 continuously increased with the increase in factor A under short priming time (less than 8.0 h), and the Gi of three other seed groups showed similar trends as those of the SVi of orhardgrass2013(Fig 4A-D).

**Fig 4.**
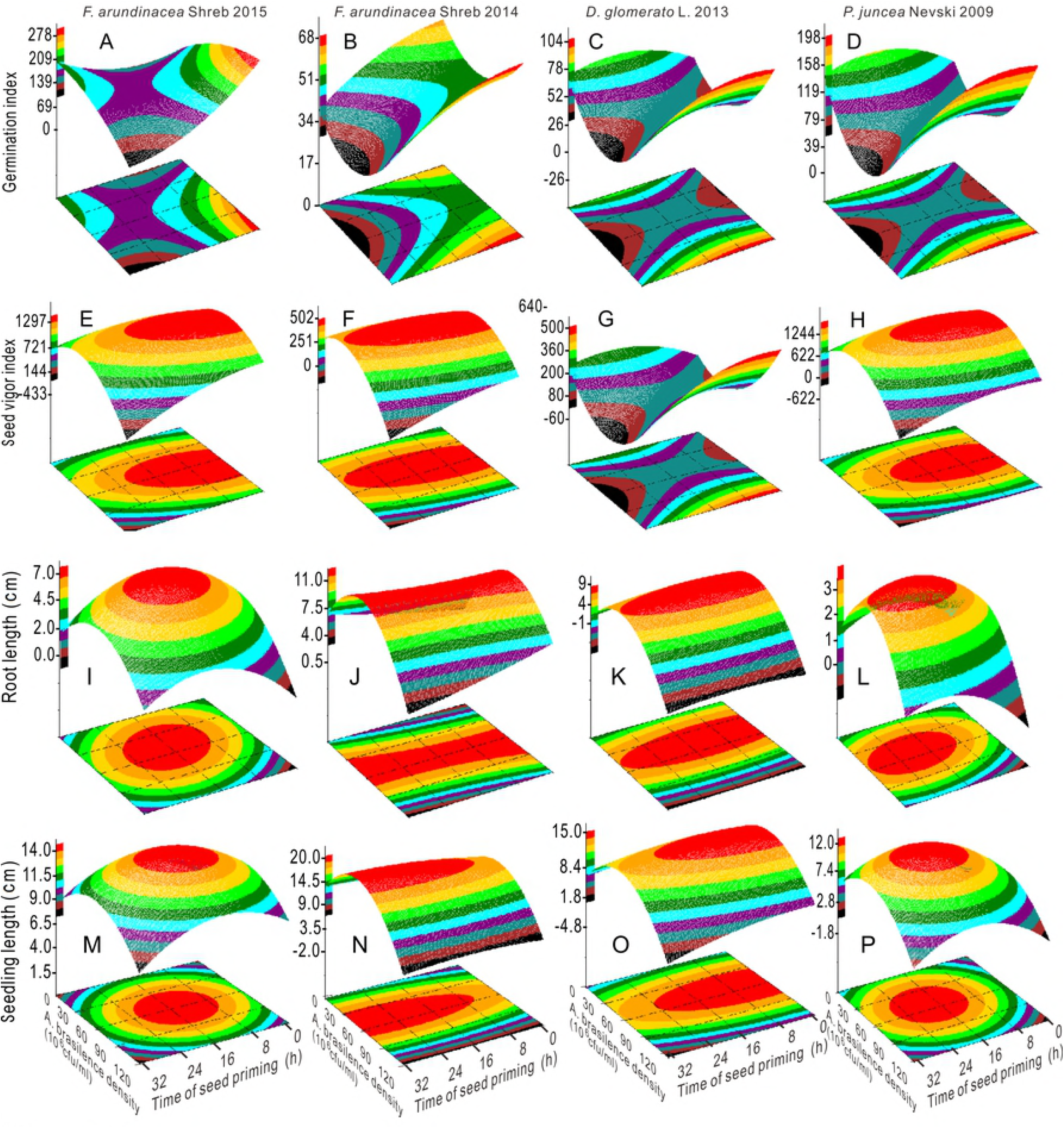
Response surface plots showing the interactions between the bacterial concentration and priming duration on the germination index (A, B, C and D), seed vigor index (E, F, G and H), root length (I, J, K and L) and seedling length (M, N, O and P).

The MDA content in Russian wildrye 2009 and the APX activity in tall fescue 2015 had minimum values at 52.8 × 10^6^ CFU mL^-1^, 19.2 h and 52.1 × 10^6^ CFU mL^-1^, 15.6 h, respectively (Figs 5B, 6F and S2 Table). The MDA content in tall fescue 2015 decreased with the increase in factor A from 60.0 × 10^6^ CFU mL^-1^ and showed opposite changing pattern as that of SOD activity. When factor A was less than 60.0 × 10^6^ CFU mL^-1^, the MDA content in tall fescue 2014 was sharply reduced with the increase in factor A (Figs 5A, C and 6D). The POD activity in Russian wildrye 2009 exhibited a parabolic trend that peaked at 61.1 × 10^6^ CFU mL^-1^ at 3.9 h; the CAT activity in Russian wildrye 2009 was positively correlated with bacterial concentration when factor A was larger than 30.0 × 10^6^ CFU mL^-1^ and when the priming time was longer than 16.0 h (Figs 5E, 6B and S2 Table). The CAT activity in tall fescue 2015 was the highest within the factor A range of 45.0-90.0 × 10^6^ CFU mL^-1^ and when the priming time was less than 4.0 h (Fig 5D). The APX and CAT activities showed analogous trends, and the APX and SOD activities were nearly complementary in orchardgrass 2013 (Figs 5F, 6C and 6E).

**Fig 5.**
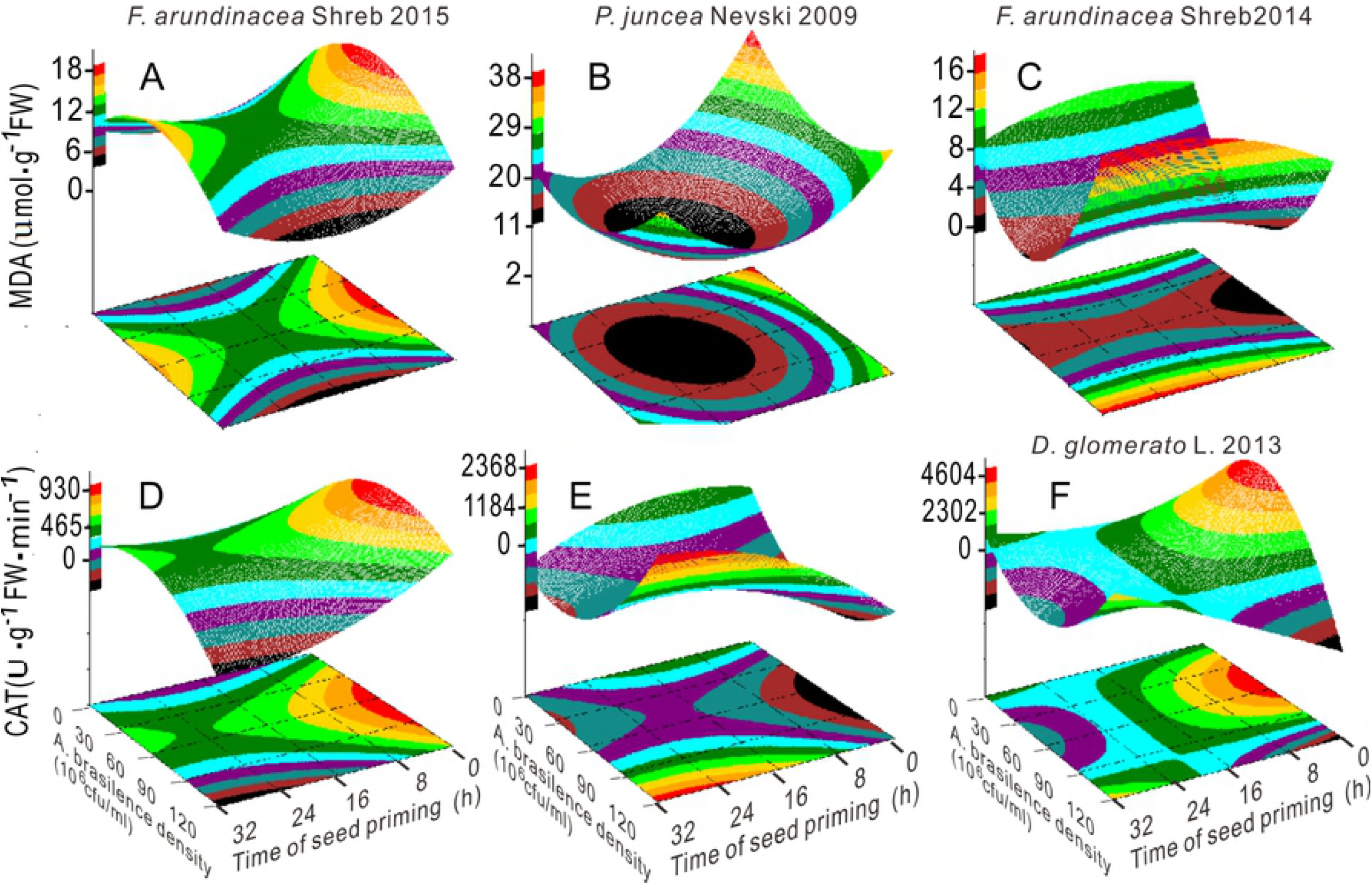
Response surface plots showing the interactions between the bacterial concentration and priming duration on the MDA content (A, B and C) and CAT activity (D, E and F).

**Fig 6.**
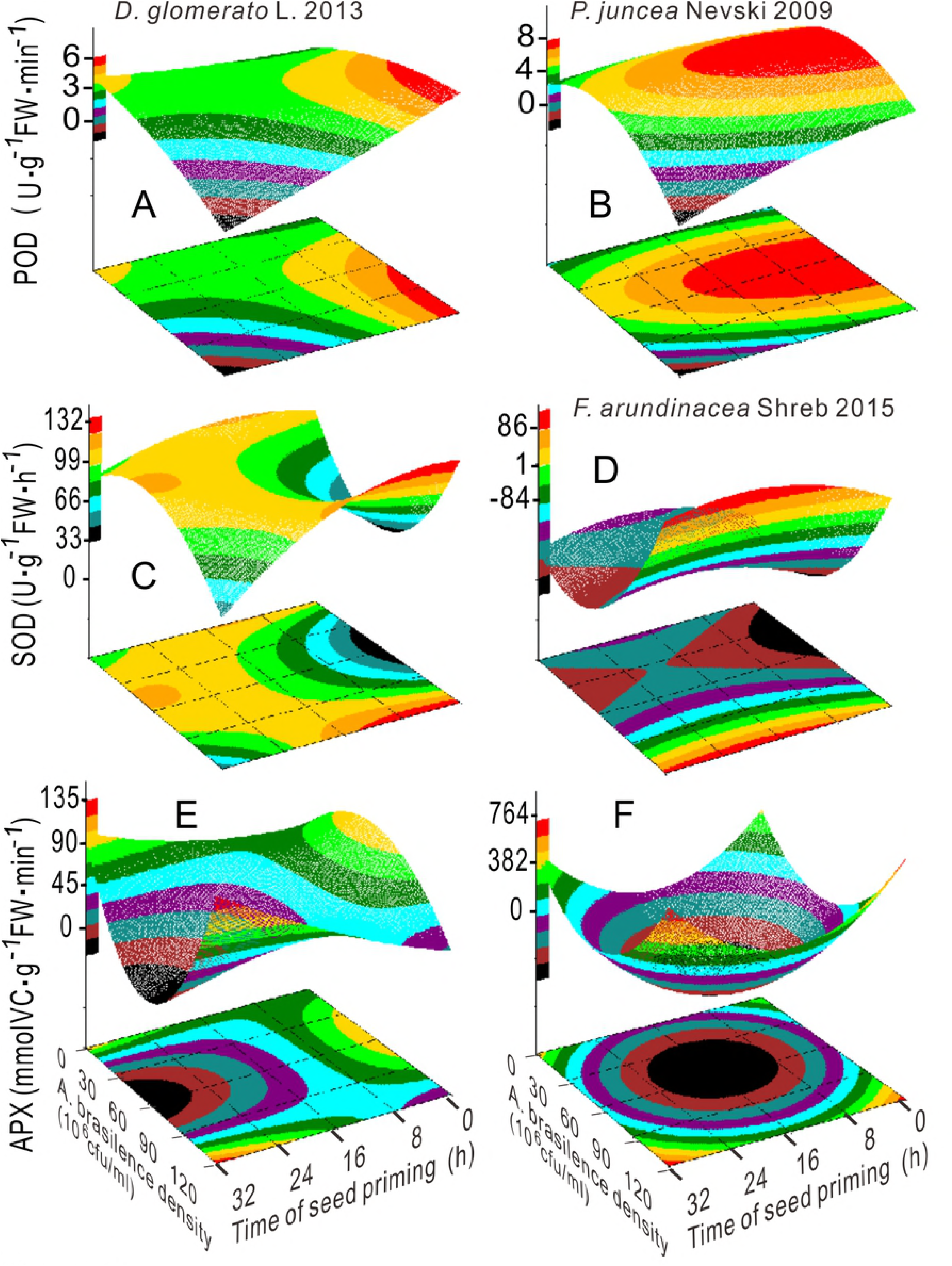
Response surface plots showing the interaction between the bacterial concentration and priming duration on the activity of POD (A, B), SOD (C, D) and APX (E, F).

A total of 48 pairs of critical values composed of factors A and C were obtained from the model analysis, as shown in Figure 7. The mode of critical values was considered the largest value and was used as the final optimal value ^[4, 25]^. The final optimal conditions for factors A and C were the bacterial concentration of *A. brasilense* of 52.3 × 10^6^ CFU mL^-1^ and the seed priming time of 17.0 h.

**Fig 7.**
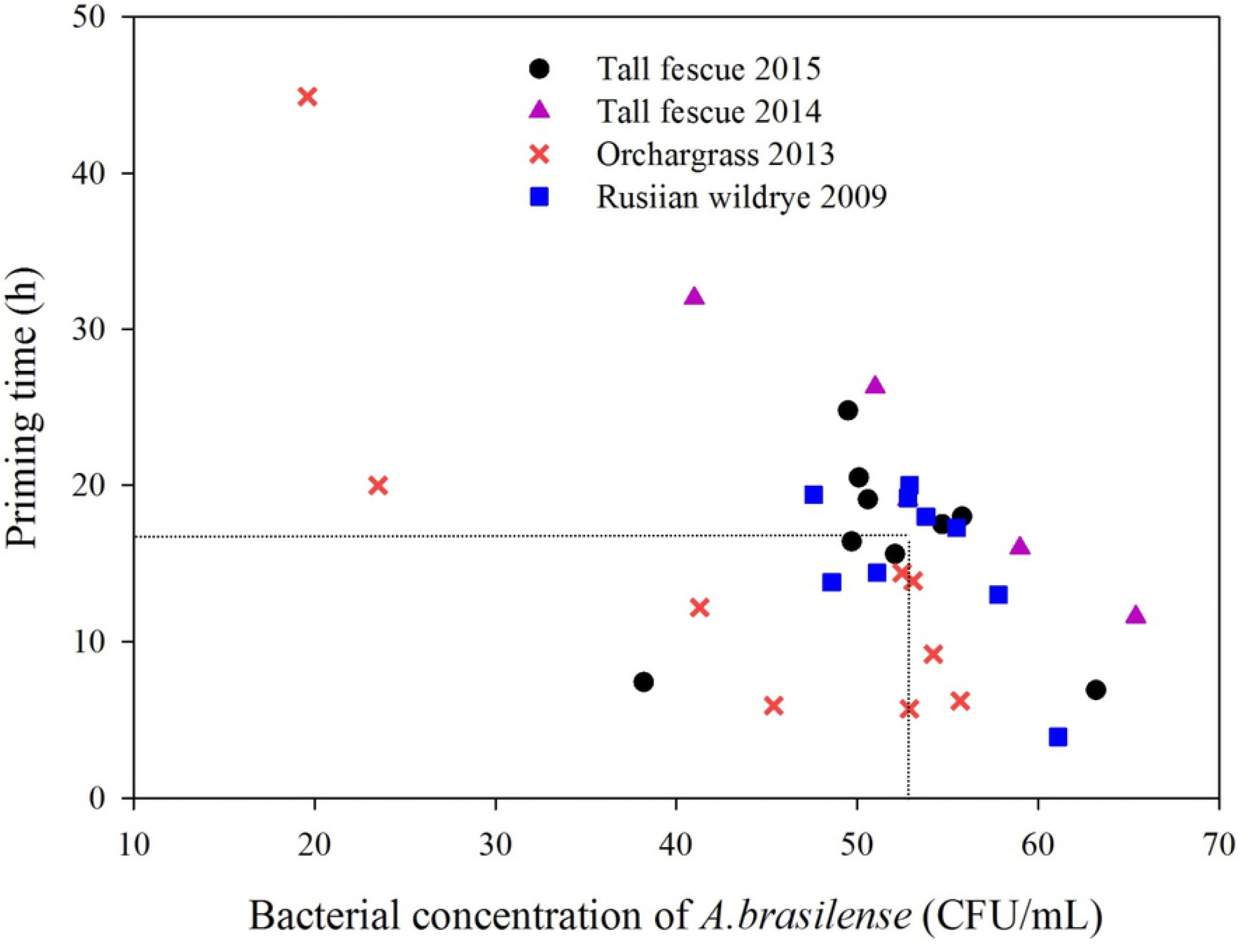
Scatter plot showing the stationary points of optimal models related to the bacterial concentration and priming duration.

## Discussion

Seed aging has become an intractable problem in agricultural science, which negatively affects the external morphology and internal physiology of seeds and seedlings ^[9]^. Seed pre-treating with PGPR is increasingly favored by scholars because of its eco-friendly and high efficiency characteristic. The present study showed that the combined treatments of *A. brasilense* inoculation and the three seed priming treatments counteracted the adverse effects from natural aging on the four groups of seeds. The results revealed that the three seed priming treatments played the dominant role in improving the seed germination and vigor. And the pairwise interactions among three factors were also significant, especially between bacterial concentration of *A. brasilense* and three seed priming treatments for the germination percentage and germination index. However, the sole function of bacterial concentration was not significant. In this regard, Carrozzi et al. noted that hydration treatment might affect only the early stages of imbibitions, while the function of *A. brasilense* is additive to the hydration process, which could result in repair actions for the promotion of aged seed germination ^[8]^. It was also found that the germination of Russian wildrye 2009 during the early phases in all treatments lagged far behind that of the other seeds. This phenomenon indicates that the germination percentage might have been intimately linked to the reparation of seed deterioration during the lag phase of germination ^[8]^. Accordingly, for the older seed, a longer lag phase was required to reverse deterioration by seed priming. Previous studies have reported that hydropriming, osmopriming and redox priming can invigorate aged seed associated with the initiation of repair- and germination-related processes, such as increased α-amylase activity, the activation and resynthesis of some ROS scavengers, the repair of chromosomal damage, protein, DNA and RNA ^[4, 26]^. On the other hand, *A. brasilense* may contribute to phytohormone communication between PGPR and aged seeds, mainly GA and IAA produced by PGPR. IAA might assist GA biosynthesis which could activate some genes responsible for α-amylase mRNA transcription in aged seeds during germination ^[27, 28]^. Therefore, we speculated here that *A. brasilense* could synergize the reinvigorating action of the three seed priming treatments for seed germination and vigor modulation in aged seeds; the latter treatments would play the main role in the early stages.

The co-treatments expressed remarkable root elongation-promoting activity, particularly under the bacterial concentration of 10^7^ CFU mL^-1^ and H_2_O_2_-priming for 18.0 h (T8) for all three aged species. The study revealed that the effect of bacterial concentration of *A. brasilense* was the greatest among the factors on root length. The root elongation response might be mainly due to the positive feedback loop between root and bacterial IAA. As reported previously, hormones produced by *A. brasilense* could affect root cell wall dynamics ^[29]^; *A. brasilense* inoculation can stimulate root carboxylate exudation acting as precursors of phytohormones, which indirectly and positively affected maize root length and area ^[30]^. In addition, under the interactive effects of factors A and C, the root and seedling lengths first increased but then decreased with the increase in factor A, there were somehow critical thresholds between the factors for the co-treatments. A moderate bacterial concentration (50.0-65.0 × 10^6^ CFU mL^-1^) and priming duration (5.8-26.3 h) could effectively facilitate the root system development in aged seeds. Similarly, *A. brasilense* Sp 7-S and Sp 245 at 10^7^ CFU mL^-1^ stimulated root and seedling growth in cucumber and lettuce ^[31]^.

To further elucidate the underlying physiological mechanisms of the combined treatments, both antioxidant enzymatic activities and lipid peroxidation were investigated. Seed aging inevitably generates the overproduction of ROS, which accelerate the disintegration of membranes. In the present study, the corresponding treatments effectively increased the activities of SOD, POD and CAT and lowered the MDA contents in the four groups of aged seeds. Especially, in severely aged seed of Russian wildrye 2009, the maximum POD activity occurred at 61.1 × 10^6^ CFU mL^-1^ and 3.9 h, and the minimum value of the MDA content occurred at 52.8 × 10^6^ CFU mL^-1^ and 19.2 h, under the strong interactive effect. It was found that the germination percentage and root length of aged seed were negatively correlated with MDA content, and were positively correlated with the SOD activity (S3 Table), which is in line with the finding of Pereira et al. ^[32]^. Wojtyla et al. pointed out that in aged seeds, the deleterious effects derived from excessive ROS can hinder cell wall loosening to allow cell elongation in root apical meristems ^[33]^. Coincidently, SOD can directly modulate the amount of ROS ^[4]^. The increased activities of antioxidant enzymes confer the ability of plants to scavenge excessive ROS ^[34]^. Therefore, reduction of MDA content and improvement of antioxidant enzyme activities in aged seeds after the co-treatment can be related to better germination and seedling growth in treated seeds compared with the untreated ones. It was clarified here that *A. brasilense* inoculation was the major modulator to improve the SOD activity in aged seeds. And this mode of action is consistent with the promoting effect for root length mentioned above. Recent studies have shown that PGPR regulate the expression of the antioxidant-responsive genes *MnSOD* and *APX1* ^[12]^. Wang et al. also indicated that a consortium of PGPR that induce drought tolerance involves SOD and *cAPX*, not POD and CAT, in cucumber ^[35]^. In this regard, the present results indicated that the three seed priming approaches were the most effective factors for enhancing POD and CAT activities. It was observed that the activities of APX and SOD were complementary in orchardgrass 2013 and that the APX activity in tall fescue 2015 had a minimum value under the interactive effects of factors A and C, which contradicts the results of some studies. However, this finding suggested that an operation mode of co-allocation might exist among antioxidant enzymes during the repair process in aged seeds. It was speculated that APX might complement the activity of SOD when the amount of ROS over-accumulated in aged seeds and that it might not play a relevant role in antioxidant enzyme system in relatively new seeds. This assumption is also supported by the study from Bosco de Oliveira et al. ^[36]^. Additionally, the reduction in MDA content after the combined treatment suggested the capability for attenuating the extent of membrane lipid peroxidation under aging stress. Although the experimental factors did not significantly affect the MDA content, the changing trends were complementary between the MDA content and SOD activity in Russian wildrye and tall fescue. Any sort of treatments reduced the MDA levels in the seedlings, which could be explained by the enhancement of antioxidant enzyme activities ^[32]^.

The combined treatments were more effective for boosting the growth of relatively aged seed (Russian wildrye 2009) compared with that of new seed. This result could confirm the repair effect of the co-treatments on aged seeds. The repair effect of co-treatments on the SOD activities, mitochondria, associated enzymes or proteins might be less/negative in new seeds with respect to old seeds, which resulted from different physiological states such as seed moisture levels, water transport and redox status ^[4]^. And seeds are expected to exhibit differences in experimental factors, which are manifested by the different genes in the different aged seeds. For tall fescue 2014, the co-treatment had a small positive impact, even negative effect on germination maybe because of its own characteristic. But it is worth noting that the germination in T2 and T7 of tall fescue 2014 occurred earlier, and T1, T2, T5, T6 and T8 significantly increased the root length, which means the co-treatment had potentially positive effect on seeds of tall fescue 2014. Therefore, the co-treatments have to be considered carefully in this species and to require further study. In addition, inter- and intra-species differences existed for the effect of factor B. The choice of seed priming treatment largely depends on plant species, variety and storage duration. The greatest average root length for tall fescue 2015, the highest average GP, Gi, and SVi for orchardgrass 2013, and those of GP, SVi, root length, shoot length, the activity of SOD and CAT for Russian wildrye 2009 occurred in the H_2_O_2_-priming. H_2_O_2_ of an appropriate level could function as a common stress and developmental signaling molecule that regulates a common set of transcription factors related to the antioxidant defense system and can eventually restore redox homeostasis and oxidative membrane damage ^[37]^. The effects of H_2_O_2_ were also correspondingly manifested by T8 (H_2_O_2_-priming) stimulating root elongation for all three aged species mentioned above. However, the germination percentage was the lowest in T9 (H_2_O_2_ priming), the germination percentage in Russian wildrye 2009 of severely aged seeds was enhanced by all nine treatments. This result implied that the level of ROS in T9 in Russian wildrye 2009 that already have an abundance of ROS, including H_2_O_2_, was not within the optimum range of concentrations, but the range was still within the “oxidative window” ^[38]^. Therefore, the H_2_O_2_ concentration lower than 10^8^ CFU mL^-1^ or priming duration shorter than 24h may be required for severely aged seeds. It is worth noting that the germination percentage of Russian wildrye 2009 in T8 was relatively higher than the remaining six treatments, and the seeds of orchardgrass 2006 germinated in T7. Taking into account the overall analysis of results of all experimental indicators, H2O2 was still the most effective condition for aged forage seeds. Similarly, H_2_O_2_ priming has improved the germination and seedling growth of pea, rice as well as aged maize, squash and tomato seeds ^[39–41]^; *A. brasilense* inoculation with H_2_O_2_-priming also enhanced the grain yield of wheat under dryland conditions ^[19]^. Therefore, *A. brasilens*e bio-priming combined with H_2_O_2_-priming constituted the optimal combination for repairing aged seeds.

Overall, a physiological mechanistic model of the combined treatments for aged grass seeds was proposed based on the current results. The PGPR bio-priming and the three seed priming treatments are coordinated but have distributed responsibilities regarding the reinvigoration of aged grass seeds during the different developmental periods. During the initiation of germination, the three seed priming treatments were the dominant factors and *A. brasilense* strains aided these activities that could lead to repair processes. For the enhancement of seedling growth (especially root length) and SOD activity, *A. brasilense* was the main driving force.

## Conclusion

*Azospirillum brasilense* inoculation and the three seed priming treatments repaired aged grass seeds by the specific operation mode of co-allocation during the different developmental stages. The three seed priming methods were the most effective factors for stimulating germination, whereas the bacterial concentration had the largest effect on root elongation. The combined treatments that elicited aging-related repair reduced the MDA content and improved antioxidant enzyme activities. Overall, *A. brasilense* bio-priming with H_2_O_2_-priming was the optimal combination for reinvigorating aged seeds. The final optimal conditions consisted of the bacterial concentration of *A. brasilense* of 52.3 × 10^6^ CFU mL^-1^ and a seed priming time of 17.0 h, given the interactive effects. However, the interactional mechanisms of combined treatments are just beginning to be unraveled. Whether the co-treatments affect aging-related proteins or genes as well as antioxidant enzyme genes still requires further exploration.

## Acknowledgements

National Key R&D Program of China (2017YFE0111000) and The National Natural Science Foundation of China (31472138) funded this work.

## Author Contributions

Q.W. and X.L. carried out the conception and designed the experiments. X.L., Z.C., W.C., Y.G., Q.L., W.Z., T.Z. and W.J. performed the experiments. Q.W., X.L., Z.C. and J.C. analyzed the data and co-wrote the manuscript. All authors reviewed the manuscript.

## Supporting information

**S1 Table. The MDA content and the activity of SOD and POD in tall fescue 2015, 2014, orchardgrass 2013 and Russian wild rye 2009.**

**S2 Table. Canonical response surface analysis for germination percentage, germination index, seed vigor index, root, shoot and seedling length, ratio of S/R, MDA content, the activity of SOD, POD, CAT and APX of the seedlings based on coded data.**

**S3 Table. Pearson correlation coefficients for activity of SOD, APX, POD, CAT, and MDA content.**

## Abbreviations

DAS: day after sowing
Gi: germination index
GP: germination percentage
RL: root length
SL: shoot length
SVi: the seed vigor index

